# An Automated Microwell Platform for Large-Scale Single Cell RNA-Seq

**DOI:** 10.1101/070193

**Authors:** Jinzhou Yuan, Peter A. Sims

## Abstract

Recent developments have enabled rapid, inexpensive RNA sequencing of thousands of individual cells from a single specimen, raising the possibility of unbiased and comprehensive expression profiling from complex tissues. Microwell arrays are a particularly attractive microfluidic platform for single cell analysis due to their scalability, cell capture efficiency, and compatibility with imaging. We report an automated microwell array platform for single cell RNA-Seq with significantly improved performance over previous implementations. We demonstrate cell capture efficiencies of >50%, compatibility with commercially available barcoded mRNA capture beads, and parallel expression profiling from thousands of individual cells. We evaluate the level of cross-contamination in our platform by both tracking fluorescent cell lysate in sealed microwells and with a human-mouse mixed species RNA-Seq experiment. Finally, we apply our system to comprehensively assess heterogeneity in gene expression of patient-derived glioma neurospheres and uncover subpopulations similar to those observed in human glioma tissue.

## Introduction

Single cell RNA-Seq is a powerful approach to quantifying cellular heterogeneity with both basic and clinical research applications^1-4^. As a result, considerable effort has been devoted to increasing the throughput and accuracy of these methods including the introduction of unique molecular identifiers (UMIs)^5^ and barcoding techniques that facilitate pooled library construction^6^. Recent advances in single cell RNA-Seq have resulted in dramatically increased scalability with a concomitant reduction in library preparation costs^7-11^. Microfluidic technology has played a crucial role in the advancement of single cell expression analysis by reducing reagent volumes, allowing high-fidelity single cell isolation, and enabling robust and automated workflows for RNA extraction and amplification^12-15^. New tools for single cell RNA-Seq exploit highly scalable microfluidic platforms, including aqueous droplets^7,8,10^ and microwell arrays^9,11^, and have facilitated miniaturization of split-pool barcoding methods for labeling cDNA libraries from hundreds or thousands of individual cells in parallel. These techniques are leading to new applications of single cell RNA-Seq including large-scale, unbiased analysis of tissues and tumors without the need for cell sorting^7^.

We recently reported single cell RNA-Seq in a solid-state microwell array platform^9^. Microwell arrays have several important advantages over droplet-based devices for single cell analysis including low sample and reagent dead volume, short cell loading time, and enhanced compatibility with short-term cell culture, cell perturbation assays, and optical imaging^16-18^. The last two features are particularly useful in minimizing sample degradation prior to cell lysis and allow the experimenter to examine and tune cell loading, identify multiplets or cell debris, and use fluorescence microscopy to determine marker composition and cell viability. In addition, high-efficiency capture of individual cells from a small sample is relatively straightforward with microwells, because cells and beads can be loaded into microwells by repeatedly flowing them over the array until all of them are captured by gravity. While our original system was capable of profiling a few hundred cells per experiment with library preparation costs of $0.10-$0.20 per cell, it suffered from several key drawbacks including low cell and molecular capture efficiency and a lack of automation^9^. Here, we report significant improvements of microwell-based single cell RNA-Seq in these three areas with no effect on overall cost. In addition, we demonstrate the compatibility of this system with the simple, 3’-end library preparation scheme SCRB-Seq^19^ and the commercially available barcoded “Drop-Seq” capture beads reported by Macosko *et al*^7^. The level of cross-contamination between wells is critically evaluated by both imaging fluorescently tracked cell lysate in oil sealed microwells and a human-mouse mixed species experiment. To demonstrate the utility of our method, we applied it to patient-derived glioma neurospheres and observed multiple phenotypic subpopulations that resemble features of intratumoral heterogeneity in glioblastoma.

## Results

### An Automated Microwell Platform for Single Cell RNA-Seq

Microwell arrays are fabricated in polydimethylsiloxame (PDMS) using standard soft lithography. Device design is highly flexible- we fabricate large arrays containing 15,000-150,000 microwells. Multiple arrays can be arranged as “lanes” on a single device for multiplexing^9^. In addition, the size of the microwells can be customized for different cell types. In the experiments described here, we use devices containing 150,000 cylindrical microwells that are 50 μm in diameter and 58 μm in height (~100 pL in volume), which accommodates most mammalian cell types. **Figure 1** illustrates the experimental work flow. New devices are first wet with a detergent-containing buffer followed by gravity-assisted cell loading. When dealing with specimens containing a small number of cells such as core biopsies or subpopulations isolated by flow sorting or laser capture microdissection, cell suspensions loaded into the device can be agitated with laminar flow to increase the fraction of cells captured by the microwell array. This process can be automated simply by connecting one end of the microwell flow cell to a standard syringe pump and reversing the flow direction repeatedly as the cells sink into the microwells (**Fig. 2A-2B**, **Supplementary Video S1**). We have used this procedure to achieve cell capture efficiencies >50%, making our system particularly useful for large-scale profiling from samples containing relatively few cells. To minimize the number of microwells containing more than one cell, we typically load <10% of the microwells. After loading the microwells, cells can be imaged with an optical microscope to assess cell viability, multiplet capture rate, cell size distribution and morphology, and surface marker composition. For mRNA capture, we load polymer beads to which barcoded oligo(dT) primers have been attached. The diameter of the beads is larger than the microwell radius, making it unlikely for more than one bead to enter a microwell and therefore facilitating high-density loading (**Fig. 2C**). After both cells and beads have been loaded into the device, another scan of the device can be performed to measure bead loading rate and the number of cell/bead pairs. The 5’-ends of the oligo(dT) primers contain a universal adapter sequence for amplification, a barcode sequence that is unique to the bead, and a second barcode sequence that for uniquely labeling a captured cDNA molecule (unique molecular identifier or UMI)^7,20,21^. The beads can capture the poly(A) tails of mature mRNA messages from eukaryotic cells and solid-phase reverse transcription results in labeling of the cDNA product with a barcode. We have demonstrated single cell RNA-Seq in PDMS microwells using capture beads that contain specific barcode sequences generated by combinatorial primer extension^9^ and, in this report, using the random barcodes sequences generated by split-pool solid phase oligonucleotide synthesis^7^. While the former have the advantage of a pre-determined sequence that can be optimized for correcting sequencing errors and other features^9^, the latter are commercially available^7^.

**Figure 1:**
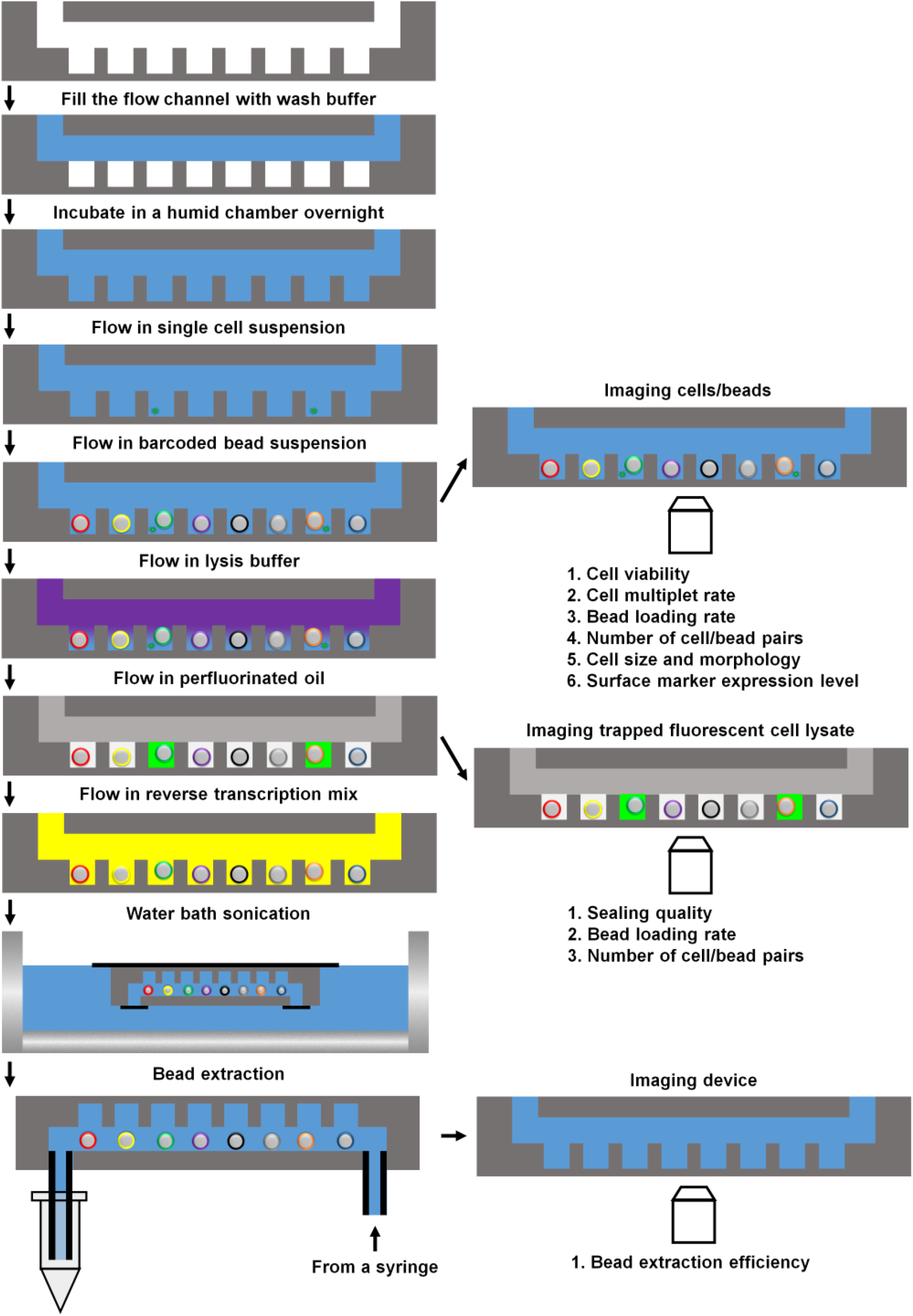
Schematic depiction (not drawn to scale) of the work flow (left column) and multiple check points (right column) of the automated microwell array platform. The check points can be used to assess the quality of a run and acquire additional phenotypic information on the same cells to be sequenced.

As described above, cell-bead pairing occurs randomly when both entities are loaded by gravity. Once this manual step is complete, cell lysis and reverse transcription occur on a computerized fluidics and temperature control system (**Fig. 2D**). We use a thermoelectric module for temperature control and an electronic, rotary selector valve to introduce different solutions to the device and reversibly seal and unseal the microwell array^9,17^. In our original report, we sealed the microwell array after introducing a lysis buffer containing a mild detergent^9^. We then used freeze-thaw cycles to initiate cell lysis, trap individual cell lysates in the microwells, and capture the liberated mRNA on barcoded beads^9^. This approach is relatively inefficient and requires low temperatures that are incompatible with automation. For efficient cell lysis, a strongly denaturing buffer that can rapidly disrupt cell membranes and deactivate nucleases would be ideal, but rapid sealing of the microwells is essential to minimize material loss and cross-contamination. Our automated system allows multiple fluids to be introduced in rapid succession, enabling the use of efficient lysis buffers without significant material loss prior to sealing. For cell lysis, we introduce a denaturing lysis buffer containing guanadinium isothiocyanate. We then rapidly introduce perfluorinated oil to seal the microwell array^22^ before cell lysis occurs. **Figure 2E** shows the lysates of isolated, fluorescently labeled cells in a microwell array following automated cell lysis and sealing. On-chip fluorescence imaging facilitates quality-control of cell viability and lysis and microwell sealing quality while providing a simple means of counting the number of cell-bead pairs and multiplet loading rate in every experiment.

**Figure 2:**
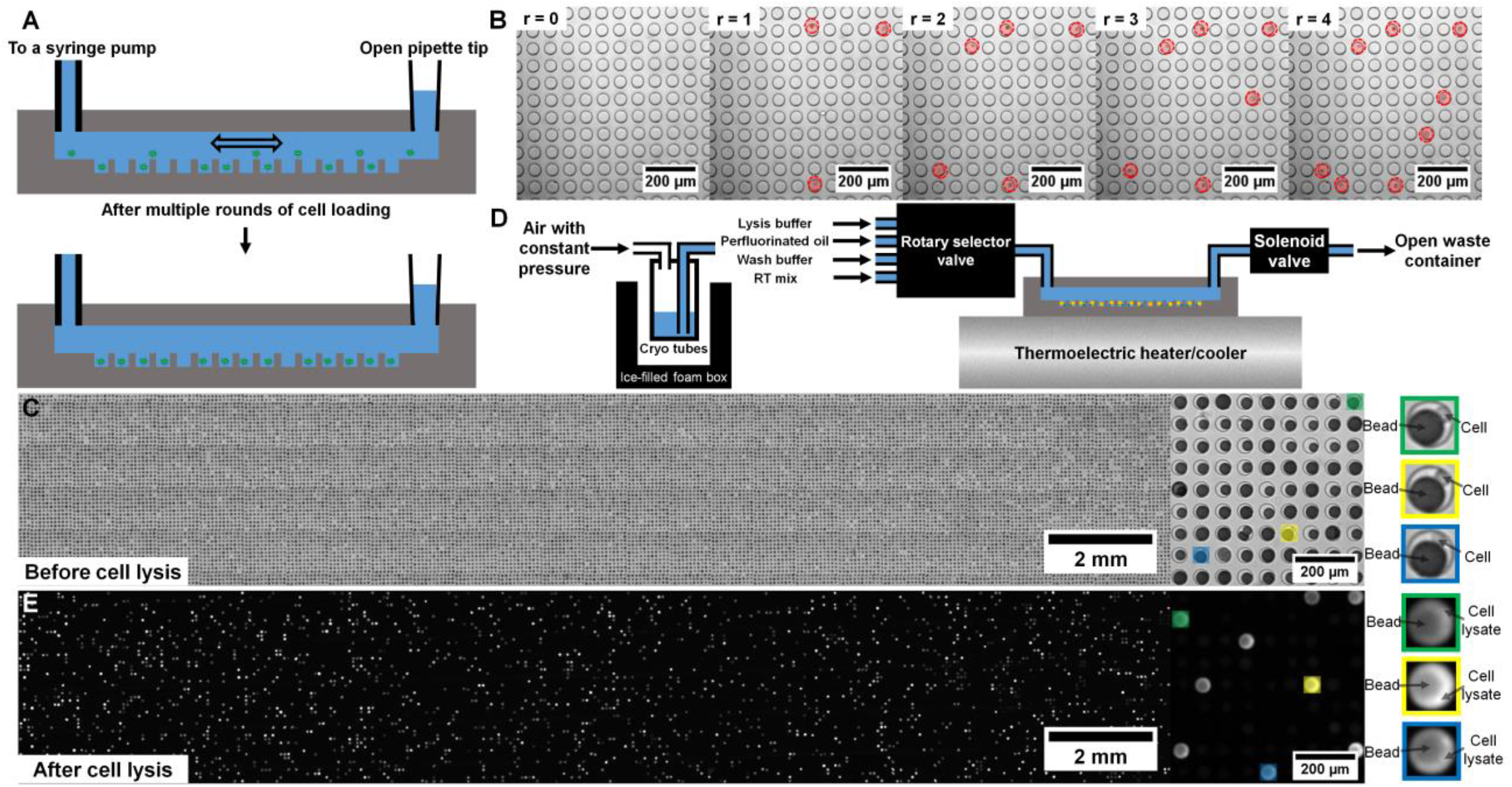
Efficient isolation of individual cells in microwells. (A) Schematic cross-section view of the device and the multi-round cell loading scheme. (B) Bright-field images of the same region of a device after several rounds (r) of cell loading. Red circles denote the loaded cells which increases monotonically with r. (C) Bright-field images of a cell/bead-loaded device. (D) Design of the automation system. (E) Fluorescence image of an oil-sealed device with trapped fluorescent cell lysate and less fluorescent bead.

Following cell lysis and mRNA capture, we introduce a detergent-containing buffer to rapidly remove the oil sealant and cell lysates. At this point the barcoded beads with hybridized mRNA are exposed to the microfluidic channel located above the microwells, and the automated system introduces all of the reagents required for reverse transcription at the appropriate temperature. Here, we use the SCRB-Seq protocol^19^ similar to what was reported for Drop-Seq^7^, and so the reverse transcription reaction also includes a template-switching step to generate full-length cDNA with universal sequence adapters on both the 3’ - and 5’-ends. Once the reverse transcription reaction is complete, we disconnect the device, remove the beads from the microwells by gentle sonication, gravity, and detergent-containing buffer flow, and complete the library construction procedure as described previously^7^. The empty device is then imaged by microscopy to measure bead extraction efficiency, which typically exceeds 99%. Note that there are still a few steps of the library construction procedure require human intervention including two PCR reactions. Further system development is required to fully automate the library construction procedure.

### High-Quality Large-scale Single Cell RNA-Seq Profiling with an Automated Microwell System

To characterize the performance of our system, we obtained RNA-Seq profiles of ~3,000 individual cells from a mixture of the human glioma cell line U87-MG and the murine fibroblast cell line NIH-3T3 in a single experiment. As shown previously, mixed species analysis is an effective approach to assessing cross-talk and purity, particularly in pooled single cell RNA-Seq experiments^7,8^. We chose the U87-MG and NIH-3T3 cell lines in order to compare the performance of our system to previous studies. We sequenced U87-MG cells in our initial report of microwell-based single cell RNA-Seq^9^ and NIH-3T3 cells were sequenced in the original report of Drop-Seq^7^.

**Fig. 3A** shows a histogram of the fraction of molecules that uniquely aligned to either the human or murine transcriptome but that aligned best to the human transcriptome for each cell. The bimodal distribution indicates that almost all of the molecules detected for roughly half of the cells originate from human mRNA versus murine mRNA for the remaining half. Because the original mixture was comprised of about 50% human and 50% murine cells, this implies that our single cell RNA-Seq profiles are quite pure (median purity of >98.8%). Cell barcodes associated with a significant number of both human and murine transcripts (<90% purity for the species with the most transcripts) likely originate from “multiplets” or instances in which two or more cells were captured in a single microwell (< 0.8% of cell barcodes).

**Figure 3:**
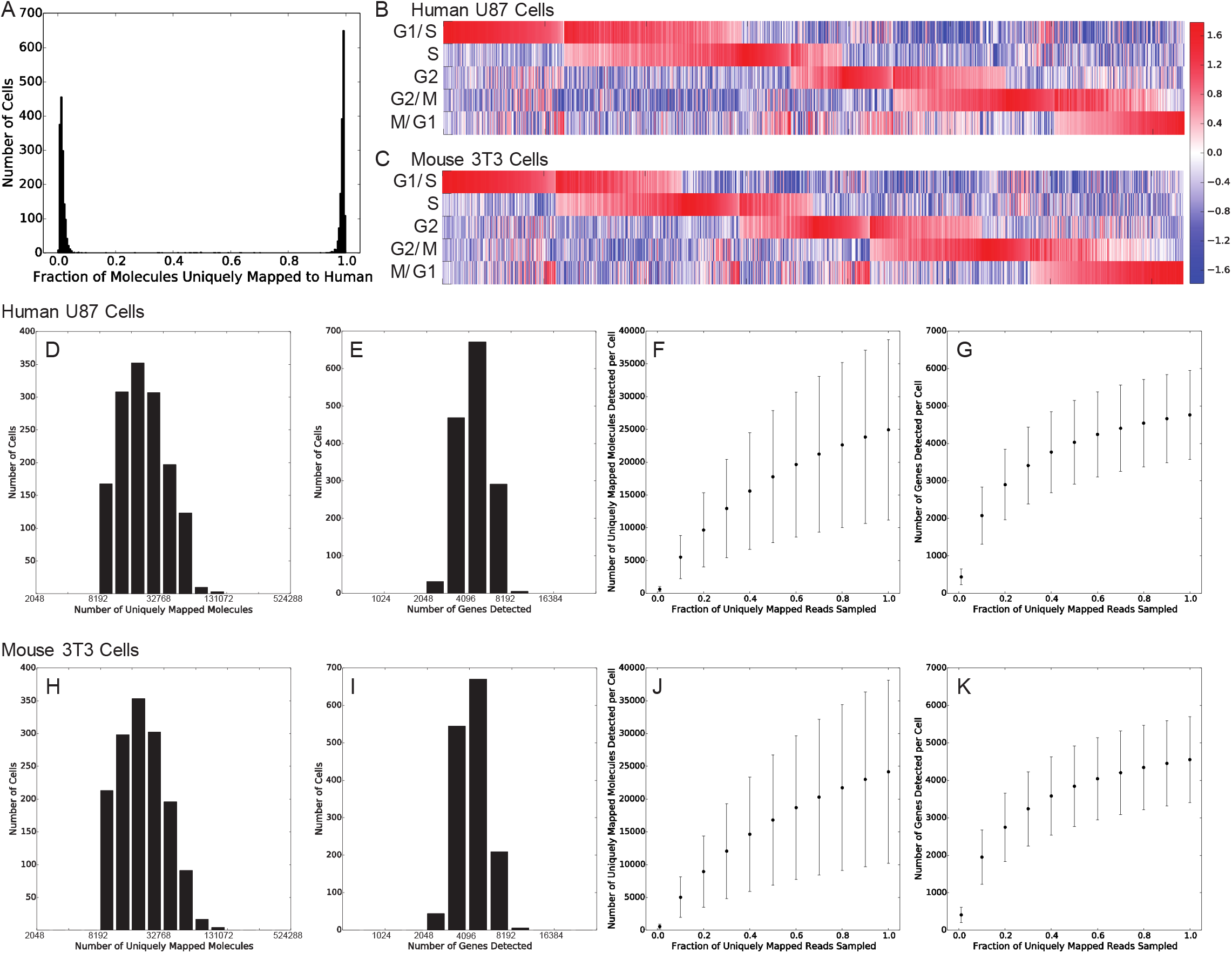
Characterization of Single Cell RNA-Seq Performance. (A) Histogram of the fraction of molecules uniquely aligned to the human transcriptome for a mixed species analysis including human U81 and murine 3T3 cells. The bimodal distribution peaked near zero and one indicate that our single cell profiles are of high purity. Cell-identifying barcodes with purities that deviate significantly from zero or one are indicative of multiplets (microwells containing two or more cells from both species). (B) Heat map of a cell cycle score for each U81 cell indicating the relative expression of genes associated with each of five cell cycle stages. The heat map shows that there are cells in all five cell cycle stages and cells that appear to be in specific intermediate transition states between stages. (C) Same as (B) but for individual 3T3 cells. (D) Histogram showing the number of uniquely aligned molecules per cell for human U81 cells. (E) Histogram showing the number of genes detected per cell for human U81 cells. (F) Sub-sampling saturation curve for the number of molecules detected per human U81 cell as a function of the number of uniquely aligned reads sampled. (G) Same as (F) for the number of genes detected per human U81 cell. (H)-(K) Same analysis as in (D)-(G) for murine 3T3 cells.

An additional indicator of purity and performance is the ability to detect subtle phenotypic subpopulations. For example, expression heterogeneity due to cell cycle asynchrony is a hallmark of single cell RNA-Seq profiles of mitotic cells. **Figs. 3B-C** show heatmaps containing cell cycle state scores for both human U87-MG cells and murine NIH-3T3 cells from this dataset (see Methods). Here, we can clearly distinguish cells in each of five stages of the cell cycle from each other as well as groups of cells transitioning between stages.

**Figs. 3D-G** show the distributions of numbers of molecules and genes detected per cell as well as saturation curves for molecule and gene detection for U87-MG cells. In our original report, we detected an average of <1,000 genes per U87 cell^9^, but here we detect ~4,800 genes per U87 cell on average. Hence, our automated microwell system has significantly higher molecular capture efficiency than our initially reported system. Similarly, **Figs. 3H-K** show the same analysis for individual murine NIH-3T3 cells. Our molecular and gene detection efficiencies are similar for the two cell lines. We detect ~25,000 molecules and ~4,600 genes per NIH-3T3 cell on average, similar to what was reported for Drop-Seq for the same cell line^7^. On average, we obtained ~208,000 raw reads per cell. Importantly, we note that neither our U87-MG nor our NIH-3T3 libraries have been sequenced to saturation. Therefore, this analysis represents an underestimate of our actual molecular and gene detection efficiencies.

We also compared our sensitivity to that of the Fluidigm C1 system for the same cell line using a publically available data set in which individual 3T3 cells were sequenced (**Supplementary Fig. S4**)^7^. Because UMIs were not implemented in these experiments, we cannot make a direct comparison of our molecular capture efficiency, but we can compare the number of genes detected per cell. We found that, at full coverage (~1 million uniquely aligned reads per cell), the Fluidigm system detected ~8,800 genes per cell on average. However, when we down-sampled the Fluidigm C1 data to ~42,000 uniquely aligned reads per cell (similar to what we obtained for 3T3 cells in this study), the Fluidigm system detected ~5,300 genes per cell. While this is comparable to the number of genes that we detected in this same cell line, the Fluidigm C1 libraries likely require more reads to reach saturation due to their full gene body coverage than our libraries in which we sequence only the 3’-end. Hence, the Fluidigm C1 library complexity and detection efficiency are most likely considerably higher than those of our platform at saturating coverage.

### Glioma Neuro-spheres Preserve Key Features of Intratumoral Heterogeneity based on Large-scale Single Cell RNA-Seq

We obtained RNA-Seq profiles of >2,200 individual cells from a patient-derived glioma neurosphere culture in a single experiment. The performance of our automated microwell array platform with these neurospheres is summarized in Supplementary Fig. S3. The mean numbers of molecules (**Supplementary Fig. S3A**) and genes (**Supplementary Fig. S3B**) detected per cell are ~14,000 and ~3,300 respectively. On average, we obtained ~303,000 raw sequencing reads per cell for TS543 cells. Saturation analysis of both the numbers of detected molecules (**Supplementary Fig. S3C**) and genes (**Supplementary Fig. S3D**) suggests that our current sequencing depth is close to saturation.

Glioma neurospheres represent an important model system for brain tumors because, in many cases, they more effectively preserve the phenotypic and genotypic features of tumors than conventional monolayer cultures^23^. They have been widely used to study drug response, glioma stem cells, and tumor progression as xenograft models^23-25^. However, to our knowledge, glioma neurospheres have not been analyzed comprehensively by single cell RNA-Seq to determine the extent of phenotypic heterogeneity and co-occurrence of cellular subpopulations within a single culture. Expression profiling of surgical specimens from glioma patients by The Cancer Genome Atlas has established classifier gene sets that stratify tumors into distinct subtypes^26^. Recent studies employing bulk expression analysis of regional heterogeneity^27^ and single cell RNA-Seq^28^ have shown that gene signatures corresponding to different patient subtypes co-occur within individual gliomas. We analyzed single cell expression profiles obtained from TS543 cells, a glioma neurosphere line that most closely resembles the Proneural glioma subtype and harbors amplification of *PDGFRA,* a genetic alteration associated with Proneural gliomas^29^. We used unsupervised dimensionality reduction and density-based cluster assignment that was uninformed of the identities of the glioma classifier genes (taken from Table S3 of Verhaak *et al.*^26^) to show that individual TS543 cells are comprised of at least two clear phenotypic subpopulations (**Fig. 4A**). For simplicity, we refer to these subpopulations as the red cluster and blue cluster. The median number of molecules detected per cell in the red and blue clusters was 11,382 and 9,771, respectively, suggesting that coverage is not a major driver of the separation between these two subpopulations. As expected, we found that Proneural genes are more commonly expressed in the majority of TS543 cells than genes from either the Classical or Mesenchymal subtypes. However, when we project expression of subtype-specific genes onto our clustering analysis, we find considerable expression heterogeneity among the classifier genes. For example, above-median expression of the Proneural classifier genes (**Fig. 4B**) is significantly enriched in the blue cluster (p < 10^-6^, hypergeometric test) whereas above-median expression of both Classical (**Fig. 4C**) and Mesenchymal (**Fig. 4D**) genes is significantly enriched in the red cluster (p < 10^-6^ for both gene sets). This phenomenon is reminiscent of the “hybrid cellular states” observed in by Patel *et al* among individual cells in human glioblastoma tissue specimens^28^. Hence, our results suggest that glioma neurosphere cultures can recapitulate the subtype-specific expression heterogeneity found in human glioma tissue.

**Figure 4:**
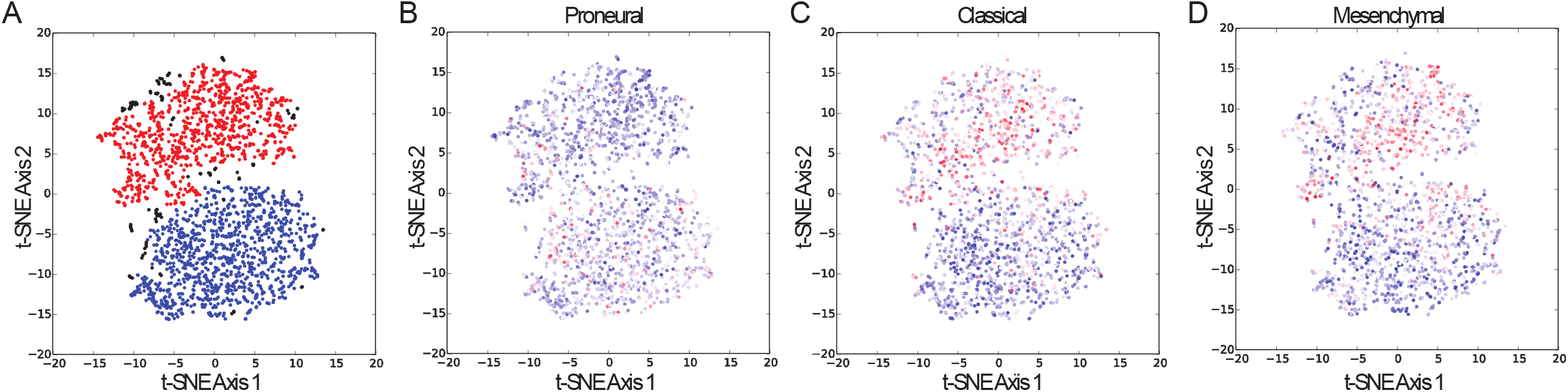
Phenotypic heterogeneity in glioma neurospheres reflects glioma patient subtypes. (A) t-SNE clustering analysis of >2,200 individual TS543 glioma neurosphere cell expression profiles showing two distinct subpopulations of cells. Assignment of individual cells to the red or blue clusters was accomplished by nearest-neighbor density analysis (see Methods). (B) Same t-SNE clustering analysis shown in (A) but colorized according expression of classifier marker genes that are characteristic of the Proneural subtype of glioblastoma (see Methods for mathematical details). Red cells have high expression of the Proneural classifier genes and blue cells have low expression of the classifier genes. (C) Same as (B) but using the classifier marker genes that are characteristic of the Classical subtype of glioblastoma. (D) Same as (B) but using classifier marker genes that are characteristic of the Mesenchymal subtype of glioblastoma.

## Discussion

We have described a significantly improved microwell platform for single cell RNA-Seq. Previous reports of similar systems suffered from several key drawbacks. The system developed by Fan *et al*^11^ was restricted to targeted analysis of specific transcripts rather than genome-wide RNA-Seq and could suffer from material loss and cross-contamination due to the lack of physical isolation between microwells. While the manual system that we reported previously was capable of unbiased RNA-Seq of individual cells, the gene detection and cell capture efficiencies were relatively low and the device had not been scaled for profiling thousands of cells in parallel^9^. Automation played an essential role in realizing the improvements demonstrated here. For example, the strongly denaturing lysis buffer employed here cannot be administered manually without significant material loss and cross contamination. An electronic fluidics system is required to introduce the buffer and rapidly seal the microwells before the cell contents escape. In addition, the Sepharose beads that we used previously for mRNA capture lack the monodispersity of the commercial mRNA capture beads used here^7,9^. Monodisperse beads of the appropriate size are essential for high efficiency cell capture because they allow us to load a single bead into most of the microwells. Mammalian cells are small enough that multiple cells can be trapped in a single microwell. Therefore, we must sparsely load the microwells with cells and rely on dense loading of the beads to maximize the frequency of cell-bead pairs in our array. The combination of monodisperse beads, the low dead volume of the microwell array device, and an efficient fluidic technique for cell loading allows us to capture >50% of cells from a suspension.

In our previously reported system, we achieved library preparation costs of ~$0.10-$0.20/cell^9^. The data set presented here included >5,000 cells from two experiments and was obtained with library preparation costs of $0.11/cell and sequencing costs of $0.48/cell. Taken together, the improvements described here have resulted in a microfluidic system for single cell RNA-Seq that is compatible with imaging and can detect thousands of genes across thousands of individual cells with a cell capture efficiency >50% and library preparation costs that are almost negligible compared to the cost of sequencing. Due to the enormous barcoding capacity of the Drop-Seq beads^7^ and the parallel fashion in which cells and beads are loaded into our prefabricated microwells, throughput of our platform, when necessary, can be further scaled up to hundreds of thousands of cells per run simply by increasing the number of microwells in a single lane and the number of lanes on a single device while keeping the time required for cell/bead loading short which is important to minimize sample degradation prior to cell lysis.

Conventional approaches to single cell analysis such as microscopy and flow cytometry are routinely employed to analyze thousands of individual cells from complex tissues. With the development of new microfluidic tools^7-11^ and an appreciation that important subpopulations can be identified with relatively shallow sequencing coverage^14^, genome-wide analysis of individual cells is beginning to reach a similar scale. As a result, new applications can be contemplated including comprehensive identification of cell types throughout an organism, simultaneous, unbiased characterization of transformed and stromal cells from solid tumors, and detection of rare cellular subpopulations that give rise to drug resistance.

## Methods

### Fabrication of PDMS Microwell Flow Cell Devices for Large-Scale Single Cell RNA-Seq

The devices are fabricated using standard SU-8 soft lithography^30^. SU-8 wafer molds are designed in Draftsight (http://www.3ds.com/products-services/draftsight-cad-software/). The diameter, height of each well, and center-to-center distance between neighbor wells are 50 μm, 58 μm and 75 μm, respectively. The height of the flow cell is 112 μm. Silanized SU-8 silicon wafer molds are obtained from FlowJEM (http://www.flowjem.com/). PDMS (Sylgard 184, Dow Corning) base and curing agent are thoroughly mixed at the ratio of 10:1, degassed under house vacuum in a desiccator (Z354074, Sigma-Aldrich) for 2 hours and is poured onto the SU-8 wafer molds in containers made of aluminum foil (01-213-100, Fisher Scientific). The degassed PDMS mixture is then cured in a 90°C oven (414004-556, VWR) for 2 hours. PDMS slabs are then gently peeled off from the molds. A 1.75 mm OD biopsy punch (15110-15, Ted Pella) is used to create inlet and outlet of flow cells. One PDMS slab with microwells and one PDMS slab with flow cell are treated in a plasma cleaner (PDC-32G, Harrick Plasma) for 30 seconds and then covalently bonded together to form the final microwell flow cell device.

### Computer-Controlled Automation for Microwell-Based Single Cell RNA-Seq

A schematic of the computer-controlled automation system is shown in **Fig. 2D**. The system consists of both temperature and fluidic control systems. Temperature control of the PDMS device is realized by directly mounting the PDMS device on top of a thermoelectric heater/cooler (CP-031, TE Technology) which is controlled through a bi-polar temperature controller (TC-36-25-RS232, TE Technology). A multi-channel selector valve (MLP777-605, IDEX Health & Science), located at the upstream of the PDMS device, is deployed to control which reservoir is connected to the device. A three-way solenoid valve (EW-01540-11, Cole-Parmer), located at the downstream of the PDMS device, is used as an on/off switch of the flow. Fluid flow is driven by a constant pressure source (3 psi) stabilized by a pressure regulator (AW20-F02, SMC Pneumatics). Because the on/off switch is located at the downstream of the device, the device is under a constant positive pressure during any incubation steps. This feature is crucial for preventing bubble formation in the device, especially at elevated temperatures such as during the reverse transcription step. To minimize dead volume, tubing with small inner diameter (127 μm, 37005T, Fisher Scientific) is used to connect reagent reservoirs and inlet of the device and that the length of the tubing is kept at minimum. This way, we are able to keep the dead volume below 10 μL which is less than the total volume of the device itself (20 to 250 μL depending on the number of wells the device has). The multi-channel selector valve is controlled by a USB digital I/O device (NI USB-6501, National Instruments). The three-way solenoid valve is controlled by the same USB digital I/O device, but through a homemade transistor-switch circuit. A C program is used to control the system.

### Library Construction and Sequencing Protocol for Microwell-Based Single Cell RNA-Seq

TS543 cell line is cultured in Complete NeuroCult™ Proliferation Medium (STEM CELL Technologies) to form neurospheres. On the day before experiment, a new device is filled with wash buffer (20 mM Tris-HCl, 50 mM NaCl, 0.1% Tween-20 (P9416-50ML, Sigma-Aldrich), pH 7.9) and stored in a humid chamber (a pipette tip box half-filled with water). On the day of experiment, TS543 neurospheres are broken into single-cell suspension by pipetting, re-suspended in TBS buffer, and stained with Calcein AM live stain dye (L3224, Thermo Fisher Scientific) at room temperature for 30 minutes. While the cells are being stained, fresh lysis buffer (1% 2-Mercaptoethanol (BP176-100, Fisher Scientific), 99% Buffer TCL (1031576, Qiagen)), RNase inhibitor doped wash buffer (0.02 U/μL SUPERaseIN (AM2696, Thermo Fisher Scientific) in wash buffer), reverse transcription (RT) reaction mix (1X Maxima RT buffer, 1 mM dNTPs, 1 U/uL SUPERaseIN, 2.5 μM template switch oligo, 10 U/uL Maxima H Minus reverse transcriptase (EP0752, Thermo Fisher Scientific), 0.1% Tween-20), and perfluorinated oil (F3556-25ML, Sigma-Aldrich) are prepared and loaded to their designated reservoirs on the automated system. Throughout the experiment, reagents in all reservoirs are chilled on ice. Cells are kept on ice once the live stain is completed. The device is then washed by TBS buffer and loaded with live-stained single cell suspension. Cells are incubated in the device for 3 minutes. Uncaptured cells are then washed away by a gentle TBS buffer wash. When few cells are available (a few to a few tens of thousands of cells), a multi-cycle cell loading scheme (**Fig. 2A-2B**) is employed to enhance cell capture efficiency. After a single cell suspension is loaded to the device, the inlet is connected to a TBS buffer-filled syringe pump (70-4501, Harvard Apparatus). The outlet of the device is connected to an open pipette tip which serves as a reservoir for overflow. The syringe pump’s operating mode is alternated between infusion and withdraw modes with a 1-minute stop time in between. We are able to achieve >50% cell capturing efficiency with 4 loading cycles in 5 minutes. Cell capture efficiency is defined as the number of cells trapped in microwells divided by the total number of starting cells in a microcentrifuge tube. The cells were counted using a fluorescence microscope (Eclipse Ti-U, Nikon). In all cases, less than 10% of the wells are loaded with cells. This minimizes the number of wells with more than one cell (multiplet loading rate). Barcoded mRNA capture beads (MACOSKO-2011-10, ChemGenes) are then loaded to the wells with the same approach. We are able to load more than 97% of the wells with beads among which less than 5% has more than one bead (**Fig. 2C**). No cell is observed in wells containing more than one bead. The device is then connected to the automated system for library preparation steps.

The library preparation work flow is adopted from Macosko *et al*.^7^ with minor modifications. Lysis buffer is first flowed through the device for 6 seconds followed by an oil flow which seals each well into a tiny (~100 pL) isolated reactor. We observed a negligible amount of cell loss after cell lysis and oil sealing steps (< 2%). The device is then incubated at 50°C for 20 minutes to enhance cell lysis and at 25°C for 90 minutes to capture mRNA onto the beads. During this period, the device is temporarily disconnected from the fluidic system, scanned on a fluorescence microscope (Eclipse Ti-U, Nikon), and then reconnected to the fluidic system. Fluorescent signal from the stained cell lysate is used to check sealing integrity. Since only a small fraction (<10%) of the wells contains a live stained cell, we expect to see the same small fraction of wells filled with fluorescent dye and that the majority of the wells to be dark if the oil sealing works well (**Fig. 2E**). In wells with a bead-cell pair, we expect to see a non-fluorescent bead surrounded by brightly fluorescent cell lysate (inserts in **Fig. 2E**). After the RNA capture step is completed, wash buffer supplemented with RNase inhibitor is flowed through the device to flush out the oil and mRNA-depleted cell lysate followed by an infusion of RT reaction mix. The device is then incubated at 25°C for 30 minutes and at 42°C for 90 minutes. At the end of the RT reaction, RNase inhibitor-doped wash buffer is used to flush out the RT reaction mix. The cDNA-coated beads are extracted from the device by a few rounds of 30-second mild water batch sonication (FS-20, Fisher Scientific) and 1-mL fast wash-buffer flow applied manually through a syringe. Bead extraction efficiency typically exceeds 99% after three rounds of sonication and wash. The extracted beads are washed once with TE/SDS buffer (10 mM Tris-HCl, 1 mM EDTA, pH 8.0), twice with TE/TW buffer (10 mM Tris-HCl, 1 mM EDTA, 0.01% Tween-20, pH 8.0), once with DI water before re-suspending in 50 μL of Exo-I reaction mix (1X Exo-I buffer, 1 U/μL Exo-I (M0293L, New England Biolabs)) at 37 °C for 30 minutes. The Exo-I treated beads are then washed once with TE/SDS buffer, twice with TE/TW buffer, once with DI water before splitting into multiple 50 μL PCR reactions (1X Hifi Hot Start Ready mix (KK2601, Kapa Biosystems), 1 μM SMRTpcr primer). We typically load about 100 cell-bead pairs per PCR reaction with 12 amplification cycles (95°C 3min, 4 cycles of (98°C 20s, 65°C 45s, 72°C 3min), 8 cycles of (98°C 20s, 61°C 20s, 12°C 3min), 12°C 5min). PCR product is purified using the solid-phase reversible immobilization (SPRI) paramagnetic bead technology (A63880, Beckman Coulter) with a 0.6:1 bead-to-sample volume ratio. Purified cDNA is then pooled together and used as input for Nextera tagmentation reactions (FC-131-1024, Illumina). We followed the standard Nextera tagmentation protocol provided by the vendor but with the following two modifications. First, 0.6 ng instead of 1 ng of cDNA is used as input per Nextera tagmentation reaction. Second, the i5 index primer is swapped with a custom primer to selectively amplify only fragments that contains the 5’ end of cDNA where cell barcodes and unique molecular identifiers (UMIs) are located. Nextera PCR product is purified in the same way as the cDNA PCR product to obtain sequencing-ready library. Representative Bioanalyzer (5067-4626, Agilent Technologies) traces of cDNA and sequencing-ready library obtained using our automated microwell platform are shown in **Supplementary Fig. S1**. The library is sequenced on a NextSeq 500 sequencer (Illumina) with 26 cycles on read 1 and 66 cycles on read 2. A custom sequencing primer is used for read 1. PhiX (FC-110-3001, Illumina) spike-in library is loaded with the single cell RNA-Seq library at 20%. We used the same primers for library construction and sequencing that were reported in Macosko *et al*^7^.

### Data Processing Procedure for Microwell-Based Single Cell RNA-Seq

Cell and molecular barcoding information are both contained in read one of our raw sequencing data, whereas all genomic information is contained in read two. We first extract the bead- and molecule-specific barcode sequences from read one. We pre-process read two using fastx_clipper by removing all poly(A) tails from the 3’-ends of each read and discarding any resulting fragment shorter than 25 nucleotides. We then map read two to either a pre-assembled human transcriptome (hg19, UCSC known genes), murine transcriptome (mm10, UCSC known genes), or a concatenated human-murine transcriptome using bwa-mem. We keep all reads with the correct strandedness that map uniquely to a specific gene with an alignment score that is greater than or equal to 85% of the length of the fragment. At this point, we associate an “address” with each non-discarded read comprised of the gene name, UMI sequence, and cell barcode sequence. In the absence of oligonucleotide synthesis, replication, or sequencing errors, each address theoretically represents a unique mRNA molecule. However, because both the cell barcode and UMI sequences are random, the address of two reads can differ simply because of sequencing errors. We first collapse all reads with the same address to a single read, keeping track of the number of reads associated with each address. Once we have assigned cell barcodes to read addresses, we examine the 8-nucletoide UMI sequences. For a given address, if there is a higher coverage address with a UMI sequence within an edit distance of one that contains the same cell barcode-gene combination, we collapse them to the same address. At this point, we consider the number of addresses to be our estimate of the number of molecules captured for each gene from each cell. We use this estimate as the basis for all subsequent analysis.

Even after the filtering procedures described above, we obtain more cell barcodes than the number of cell-bead pairs loaded in our device. These additional cell barcodes arise from several sources including additional sequencing or synthesizer errors and the beads that are not paired with a cell, which can capture low levels of ambient RNA during the experiment. Nonetheless, as shown in **Supplementary Fig. S2**, we can readily identify a population of very high coverage barcodes based on the distribution of captured molecules that is consistent with the number of cell-bead pairs imaged in our device. Similar observations have been made in previous studies^7,9^.

### Cell Cycle Analysis

We adopted the cell cycle analysis method developed by Macosko *et al*^7^. Please refer to the original paper for details. Briefly, the expression level of a set of genes that are known to reflect different phases of cell cycle were used to calculate a phase-specific score for each cell. Each cell is then classified into one of the ten patterns of phase-specific scores (including eight potential patterns along the cell cycle and two patterns for equal scores of all phases (either all active or all inactive)) based on the maximal correlation of the cell’s phase-specific score with these ten patterns. Cells within each class were further ordered based on their relative correlation with the preceding and succeeding patterns. The set of genes used to calculate the phase-specific scores were obtained from the Supplemental Figure 15 in Whitfield *et al* which reflect five phases of cell cycle (G1/S, S, G2, G2/M, M/G1)^31^. The eight potential patterns along the cell cycle that the cells were classified into are: only G1/S is on, both G1/S and S are on, only S is on, both S and G2 are on, only G2 is on, both G2 and G2/M are on, only G2/M is on, both G2/M and M/G1 are on.

### Clustering Analysis of Single Cell Expression Profiles

We clustered our TS543 single cell expression profiles using a set of highly variable genes identified based on a dispersion analysis of the entire data set. We first normalized the molecular counts for each gene in each cell by the total number of molecules detected in that cell. We considered these normalized molecular counts to be expression levels. Next, we plotted the coefficient of variation vs. mean expression across all genes detected in at least five cells and grouped the genes into 50 evenly-spaced bins based on log-transformed expression levels. We computed a z-score for each bin and took genes with a z-score greater than three to be highly variable given their expression levels as long as they were detected in at least 10% of cells (see **Supplementary Table S1** for a complete list). Hence, the variance in these genes is less likely to result from technical noise and more likely to result from real biological variation. We then computed a matrix of Pearson correlation coefficients between the log-transformed expression profiles of each cell using only the highly variable genes. Finally, we used this Pearson correlation matrix as input to the t-stochastic neighborhood embedding (t-SNE) algorithm^32^ for unsupervised clustering as implemented in the Python scikit-learn package. The results of the t-SNE clustering are displayed in **Fig. 4**. We assigned cells to discrete clusters by density analysis with the DBSCAN function in scikit-learn using the Euclidean distance metric.

We used the following score, *S_subtype,i_*, to assess expression of glioma subtype-specific genes in an individual cell *i*:

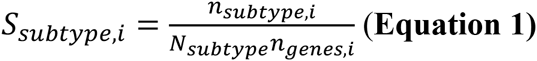

where *n_subtype,i_* is the number of subtype-specific genes detected in cell *i*, *N_subtype_* is the number of subtype-specific genes detected in the entire dataset, and *n_genes,i_* is the number of genes detected in cell i.

### Analysis of Single Cell RNA-Seq Data Generated by the Fluidigm C1 System

As described above, we sequenced NIH-3T3 murine fibroblasts as part of a performance test for our system. This same cell line was sequenced using the Fluidigm C1 system by Macoscko *et al*^7^. We downloaded the raw SRA data for these experiments from GEO accession GSE701151 and converted these data to 192 fastq files, corresponding to 192 single cell profiles using fastq-dump in the SRA Toolkit package. We then aligned the each fastq file to a concatenated human-mouse pre-assembled transcriptome using bwa-mem and identified uniquely aligned reads just as described above. Because the Fluidigm C1 data set originated from a mixed species experiment in which human HEK cells were mixed with murine 3T3 cells, we identified cells with >90% of the reads aligned to the murine transcriptome and quantified the number of genes detected per cell at two different read depths (**Supplementary Figure S4**).

## Acknowledgements

The authors thank Dr. Sohani Das Sharma for assistance with cell culture and preparation, Erin C. Bush for assistance with library preparation and sequencing, and Dr. Harris Wang for the loan of a syringe pump. P.A.S. is supported by K01EB016071 from NIH/NIBIB, R33CA202827 from NIH/NCI, and U54CA193313 from NIH/NCI.

## Author Contributions

J.Y. and P.A.S. conceived and designed the automated microwell array system. J.Y. fabricated the microwell array devices, constructed the automated system, implemented the library construction protocol, and generated the single cell RNA-Seq data. J.Y. and P.A.S. analyzed the data and wrote the paper.

## Accession Codes

The RNA-Seq data generated in this study has been deposited in the Gene Expression Omnibus hosted by the National Center for Biotechnology Information under accession GSE85575.

## Additional Information

Competing Financial Interests Statement: Columbia University has filed a patent application based on this work.

